# Urinary candidate biomarkers in an experimental autoimmune myocarditis rat model

**DOI:** 10.1101/191379

**Authors:** Mindi Zhao, Jiangqiang Wu, Xundou Li, Youhe Gao

## Abstract

Urine is a better source than plasma for biomarker studies, as it can accumulate all changes in the body. Various candidate urinary biomarkers of physiological conditions, kidney disease and even brain dysfunction, have been detected in urine; however, urine has rarely been used to reflect cardiac diseases. As the clinical presentations of myocarditis are heterogeneous, reliable and sensitive diagnostic biomarkers of myocarditis are very important. In this study, candidate urinary biomarkers in the myosin-induced autoimmune myocarditis rat models were characterized using the isobaric tandem mass tag labeling approach coupled with high-resolution mass spectrometry. Compared with controls, forty-six urinary proteins were significantly changed in the myocarditis rats; among them, ten had previously been associated with myocarditis, twelve corresponding gene products have been annotated as mainly cardiovascular network genes by the Ingenuity Pathway Analysis, and four urinary proteins were validated by western blot.

## Introduction

As urine can accumulate all types of changes in the body, it is a good source for biomarker studies [1]. Candidate biomarkers of various diseases have been detected in urine, and some of them even perform better in urine than in plasma [2]. In addition, major advances in the identification of urine proteins allow them to play an important role in biomarker discovery [3]. Previous studies have shown that kidney diseases and even brain diseases can be reflected in urine [4, 5]; therefore, the present study sought to determine if other diseases can be monitored in urine.

Myocarditis is characterized by inflammation of the myocardium and is triggered by various situations, including autoimmunity and infection[6]. Myocarditis is one of the leading causes of sudden cardiac death in young adults[7] and athletes[8] and can progress into potentially devastating sequelae, such as dilated cardiomyopathy (DCM)[9]. Therefore, an early and reliable diagnosis of myocarditis is very important. However, the clinical presentations of myocarditis are heterogeneous, ranging from asymptomatic infection to fatigue, chest pain, arrhythmias and even heart failure, which increase the difficulty of diagnosis. Endomyocardial biopsy is the gold standard in the evaluation of myocardial status, but it is not commonly used in routine clinical practice[10]. Cardiovascular Magnetic Resonance (CMR) is a good noninvasive tool for the diagnosis of acute myocarditis, but its sensitivity is susceptible to other factors[11]. The predictive values of some cardiac biomarkers, such as CK or CK-MB, are too low to be used in clinical screening[12]. However, troponin T or I perform better than CK or CK-MB and can help confirm the diagnosis. The specificity of troponin I or T is very high, whereas its sensitivity is relatively low[13, 14]. Therefore, none of these serum biomarkers have been shown to be an effective tool for accurately predicting myocarditis, and new diagnostic methods and biomarkers are urgently needed to improve the identification of myocarditis.

Animal models are a good tool for the study of disease urinary biomarkers, as the exact start of the disease is known and there is very few confounding factors. Myosin-induced autoimmune myocarditis (EAM) is a model of inflammatory heart disease that is mediated by various types of immune cells and their products[15]. As the initiators and mediators of disease progression, T cells play important roles in EAM modeling. The myocardium of patients with myocarditis and DCM are infiltrated with CD4+ and CD8+ T cells[16], which is also reflected in the EAM model. The histopathology of rodents immunized with cardiac myosin demonstrates severe myocarditis [17]. As a result, the EAM model has tremendously improved our understanding of the molecular and immunological mechanisms of myocarditis and DCM [18].

In this study, we sought to characterize the urinary biomarkers of myocarditis. EAM-induced rats were used to model the progression of the disease (Figure 1). Changes in the urinary proteins were identified by the isobaric tandem mass tag (TMT) labeling approach coupled with high-resolution mass spectrometry (LC/MC) to enable an in-depth coverage.

**Figure 1.**
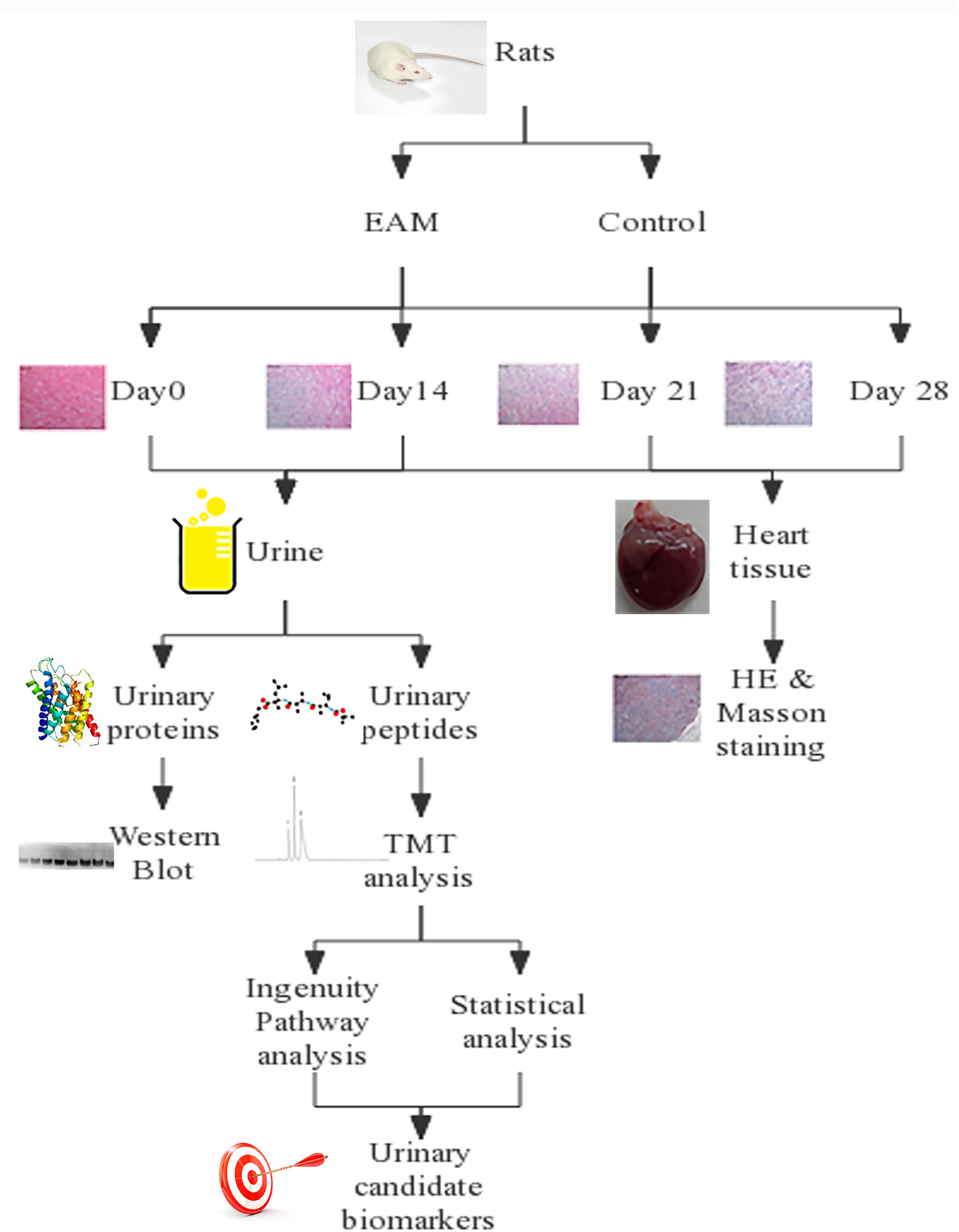
The workflow of urinary proteome analysis in EAM rats. Urine was collected at day 0, 14, 21 and 28. After urine collection in each phase, histological and morphometric analyses of the heart were conducted. Urinary proteins were identified by TMT-labeled LC/MS/MS.

## Materials and methods

### Experimental rats

Thirty male Lewis rats (8 weeks old) were purchased from the Institute of Laboratory Animal Science, Chinese Academy of Medical Science & Peking Union Medical College. The experiment was approved by the Institute of Basic Medical Sciences Animal Ethics Committee, Peking Union Medical College (Animal Welfare Assurance Number: ACUC-A02-2014-007). The study was performed according to guidelines developed by the Institutional Animal Care and Use Committee of Peking Union Medical College.

### Experimental autoimmune myocarditis rats

EAM was induced in Lewis rats as previously described[19]. All rats (n = 30) were randomly divided into two groups. All rats were anesthetized on days 0 and 7; rats in the EAM group were immunized and intradermically injected with 1 mg of porcine cardiac myosin (Sigma, St. Louis, MO) emulsified in complete Freund’s adjuvant (CFA). Rats in the control group were administered CFA and infused with an equal amount of saline. On days 0, 14, 21 and 28, all rats were individually placed into metabolic cages for four hours to collect urine. At each time point, three rats in the EAM group and the control group were sacrificed after urine collection, and their hearts were collected for histological and morphometric analysis. Body weight was measured daily.

### Histological analysis

The hearts of EAM and control rats were harvested 0, 14, 21 and 28 days after immunization. For histopathology, the heart was fixed in formalin (4%) and embedded in paraffin. Paraffin-embedded heart sections were stained by Masson’s trichrome staining to reveal fibrosis. The histopathological lesions were evaluated with hematoxylin and eosin (HE) staining.

### Urine sample preparation

Urine from the EAM and control groups was centrifuged at 2,000 g after collection. After removing the pellets, three volumes of acetone were added. After centrifugation, lysis buffer (8 M urea, 2 M thiourea, 25 mM dithiothreitol and 50 mM Tris) was used to dissolve the pellets. Then, proteins were digested with trypsin (Promega, USA) using filter-aided sample preparation methods [12]. Briefly, proteins were denatured by dithiothreitol and alkylated by iodoacetamide. Proteins were then digested with trypsin (1:50) at 37°C overnight. The digested peptides were desalted using Oasis HLB cartridges (Waters, USA).

Six urine samples from the EAM and control groups on day 21 were individually labeled with 126, 127, 128, 129, 130 and 131 TMT reagents according to the manufacturer’s protocol (Thermo Fisher Scientific, Germany) and then analyzed with two dimensional LC/MS/MS.

### Reverse-phase liquid chromatography (RPLC)

All samples were fractionated using offline high-pH RPLC columns (XBridge, C18, 3.5 μm, 4.6 mm × 250 mm, Waters, USA). The samples were diluted in buffer A1 (10 mM NH_4_FA in H_2_O, pH = 10) and then loaded onto the column. The elution gradient consisted of 5–30% buffer B1 (10 mM NH_4_FA in 90% acetonitrile, pH = 10; flow rate = 1 mL/min) for 60 min. The eluted peptides were collected at one fraction per minute. After lyophilization, the 60 fractions were resuspended in 0.1% formic acid and concatenated into 30 fractions by combining fractions 1 with 31, 2 with 32, and so on.

### LC-MS/MS analysis

Each fraction was analyzed duplicate using a reverse-phase C18 (3 μm, Dr. Maisch, Germany) self-packed capillary LC column (75 μm × 120 mm). The eluted gradient was 5%–30% buffer B (0.1% formic acid in acetonitrile; flow rate 0.3μl/min) for 60 min. The peptides were analyzed using a TripleTOF 5600 system. The MS data were acquired in data-dependent acquisition mode using the following parameters: 30 data-dependent MS/MS scans per full scan; full scans were acquired at a resolution of 40,000 and MS/MS scans were acquired at 20,000; rolling collision energy; charge state screening (including precursors with +2 to +4 charge state); dynamic exclusion (exclusion duration 15 s); MS/MS scan range of 250-1800 m/z; and scan time of 50 ms.

### Data processing

All MS/MS spectra were analyzed by the Mascot search engine (version 2.4.1, Matrix Science, UK), and proteins were searched against the Swissprot_2014_07 databases (taxonomy: Rattus, containing 7,906 sequences). The parameters were set as follows: the carbamidomethylation of cysteines and TMT labeling was set as fixed modifications; two missed trypsin cleavage sites were allowed; the precursor mass tolerance was set to 10 ppm; and the fragment mass tolerance was set to 0.05 Da.

Proteins were filtered using the decoy database method in Scaffold (version 4.3.2, Proteome Software Inc., Portland, OR). Protein identifications that were greater than 99.0% probability and achieved a false discovery rate (FDR) less than 1.0% were allowed [14]. Each protein contained at least 2 unique peptides. Protein probabilities were assigned using the Protein Prophet algorithm.

Scaffold Q+ was employed for the quantification of TMT labeling. The parameters were set as follows: the statistical test used in Scaffold Q+ was the permutation test; the calculation type was the median; the reference type was average protein reference; and the normalization between samples was set to “ON”.

### Western blot

Primary antibodies against regenerating islet-derived protein 3 (REG3), myoglobin, cystathionine gamma-lyase (CSE) and beta 2-microglobulin (B2M) (Abcam, Cambridge, U.K.) were used to validate the MS results. A total of 30 μg of urinary proteins from the EAM and control groups on days 0, 14 and 21 (three samples per time point) was separated by SDS-PAGE and transferred to a polyvinylidene fluoride (PVDF) membrane (Whatman, UK). The membranes were blocked in 5% milk and then incubated with primary antibodies overnight. After washing, the membranes were incubated with peroxidase-conjugated anti-rabbit IgG (ZSGB-bio, China). Finally, the proteins were visualized using enhanced chemiluminescence reagents (Thermo Fisher Scientific, USA). ImageJ analysis software (National Institutes of Health, USA) was used to analyze the intensity of each band.

### Functional analysis

Ingenuity Pathway Analysis (IPA) was used to analyze the functional annotation of differential proteins between the EAM and control groups. The differential proteins were uploaded onto the IPA website and then compared to information from the curated literature. The biological functions assigned to each canonical pathway were ranked according to the significance of that biological function to the pathway. All identified proteins were also analyzed by IPA software to enable a detailed annotation, including cellular components, metabolic processes and canonical pathways.

## Results and discussion

### Body weight and histopathological characterization over time

In the EAM group, all fifteen rats developed myocarditis, and none of them died before they were sacrificed. In the control group, all rats appeared normal after the administration of CFA and infusion with equal amounts of saline. On day 0, the mean body weight (BW) of the EAM and control rats was 204.0 g (±5.6 g) and 212.6 g (±11.4 g), respectively; on day 28, the mean BW of the EAM and control rats was 329.8 g (±13.9 g) and 326.5 g (±13.2 g), respectively. There was no significant difference in BW between the two groups at any time after immunization.

Macroscopic evaluation of the whole heart showed that the hearts maintained a normal shape between days 0 and 14. On day 21, a small discolored area was observed in the hearts of EAM rats, and representative heart sections from this area were then used for the histopathological examinations. On day 28, the discolored area was enlarged, and some areas were scarred. The scarred areas were then used for histopathological characterization.

Histopathological examinations of the hearts of EAM rats were performed to determine the severity of the disease process. On day 14 after the first immunization, no pathological changes were observed in the heart, as hematoxylin-eosin (H&E) staining of the EAM heart appeared similar to that of the control hearts. On day 21, H&E staining of the EAM heart revealed the presence of numerous mononuclear inflammatory cells, which is consistent with previous studies[20, 21]. On day 28, large areas in the H&E-stained EAM hearts displayed inflammatory cellular infiltrate; there were also more necrotic swollen myocytes in the EAM hearts than in the control hearts, and many myocardial fibers were destroyed in the EAM hearts (Figure 2A).

**Figure 2.**
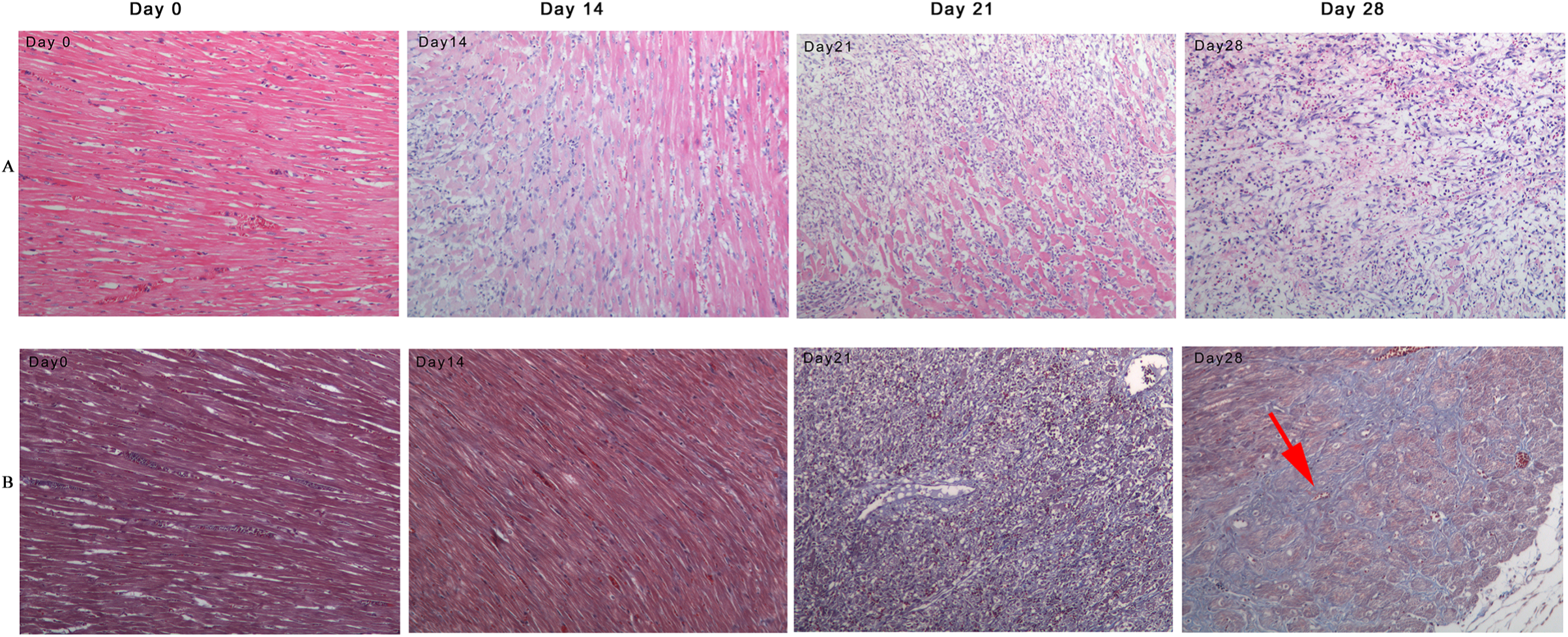
Histopathological characterization 0, 14, 21 and 28 days after immunization. A) H&E staining of the EAM heart. B) Masson’s trichrome staining of the EAM heart; the red arrow indicates collagen deposition.

The heart tissues of these EAM rats were also analyzed by Masson’s trichrome staining to demonstrate the accumulation of extracellular collagenous matrix. As with the H&E staining, on day 14, the EAM hearts did not display obvious inflammatory filtration. However, on day 21, the myocytes were partially destroyed in some sections of the EAM hearts. On day 28, the EAM heart sections stained by Masson’s trichrome showed obvious collagen deposition (Figure 2B). These pathological changes revealed the successful induction of heart failure in the EAM rats.

### Changes in urinary proteins between the EAM and control rats

Macroscopic and microscopic evaluation of the heart revealed that on day 21, inflammatory cell infiltration was first detected in a few sections, and on day 28, the number of sections displaying inflammatory cell infiltration increased remarkably. Urine samples from day 21 were chosen for MS analysis based on preliminary studies, which showed that some sections of EAM rat hearts began to exhibit inflammatory cell infiltration at this time point; thus, the changes in urinary protein on day 21 may be specifically related to the inflammatory response. To improve the accuracy of quantitative identification and to increase protein coverage, the analyzed peptides were labeled by TMT agents and then characterized in duplicate using offline high RPLC coupled with high-resolution MS. All raw spectra were searched against Swissprot by Mascot software and then filtered and quantitated by Scaffold software. Proteins that could not be differentiated by MS/MS spectra were reported as one protein group. The workflow in Figure 1 provides a brief overview of the processes and technical design of this study.

A total of 851 proteins was identified at a protein FDR of 1% and included at least two unique peptides. It was a thorough characterization of the rat urine proteome. Among them, 816 proteins were quantitated by TMT-labeling analysis in both technical replicates (Supplemental Table 1), implying that most proteins can be identified and quantitated in each urine sample. Quantitative proteomics analysis by Scaffold Q+ provided insights into changes in the protein profile of rat urine due to myosin-induced EAM. After calculation by the software, fold changes of all urinary proteins in each rat could be exported.

**Table 1.**
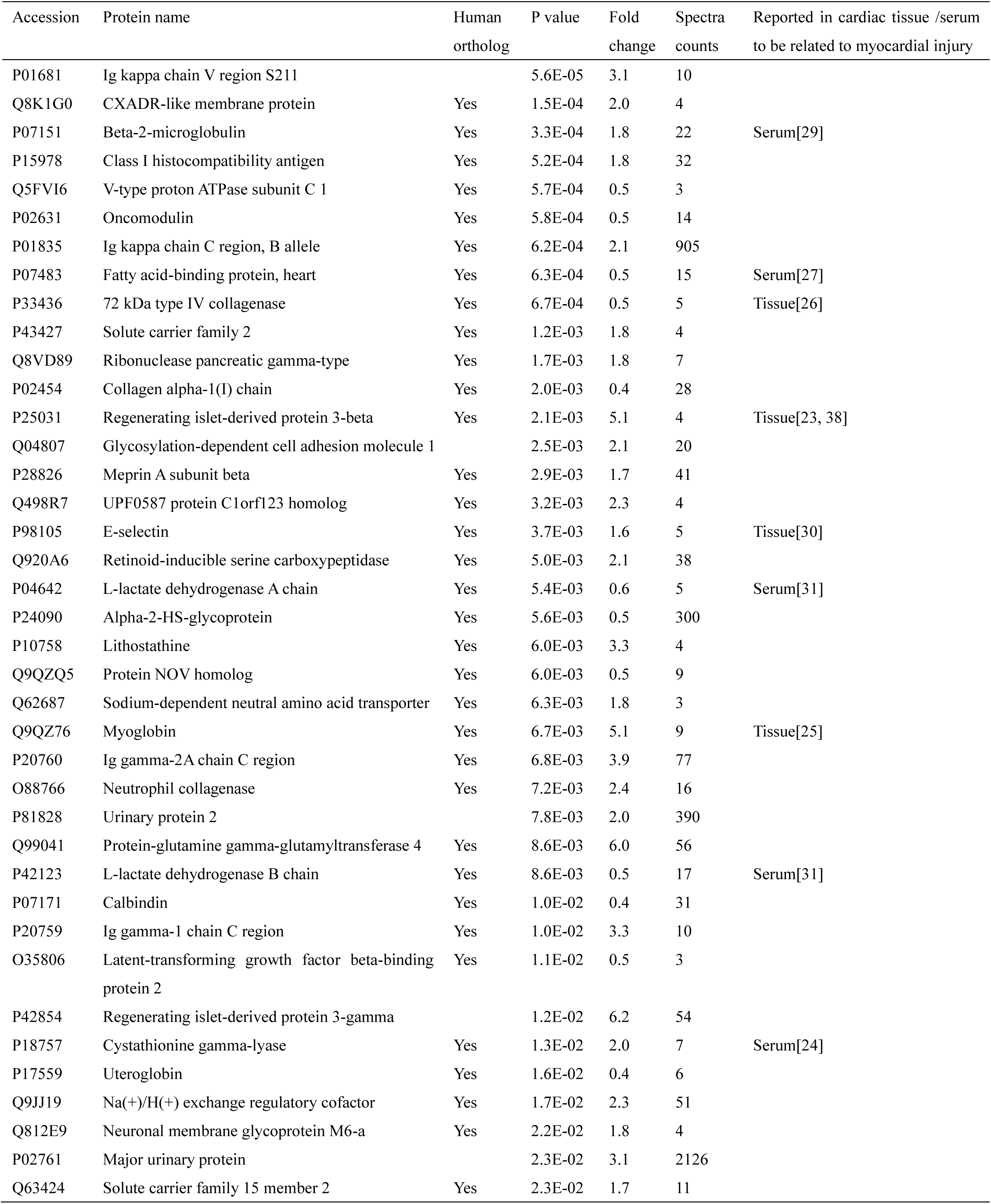
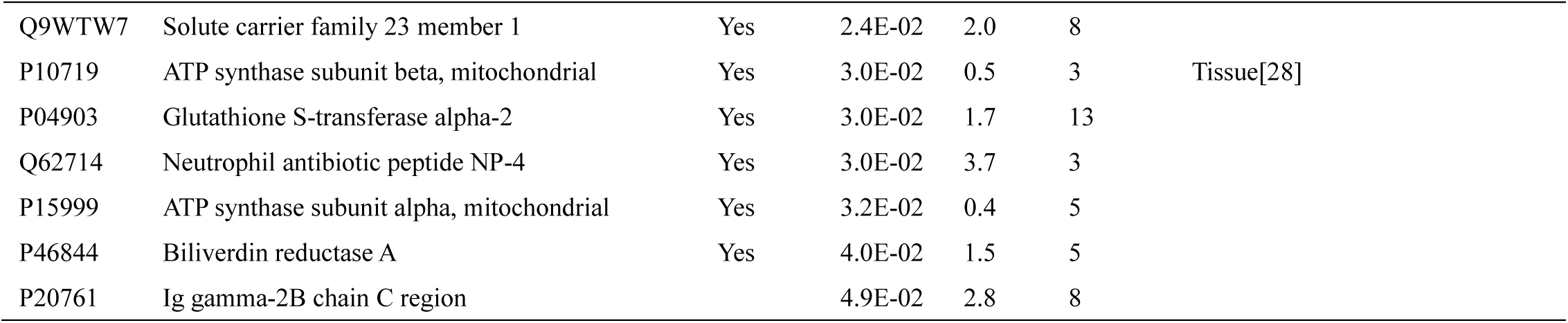
Changed urinary proteins in EAM rats.

Unsupervised clustering of the proteomic profiles of 816 matched proteins from the 6 urine samples (each with a technical replicate) from day 21 was performed (Figure 3C). A heat map was constructed based on the quantitative data, which were extracted from the Scaffold software. Clustering analysis of the quantitative data showed that replicates from the same sample were most closely clustered together (except for E2 & E3). As shown in the figure, the proteomic profiles within the same group were similar to each other and distinctively different from the other group. Varying differences between the control and EAM groups were revealed.

**Figure 3.**
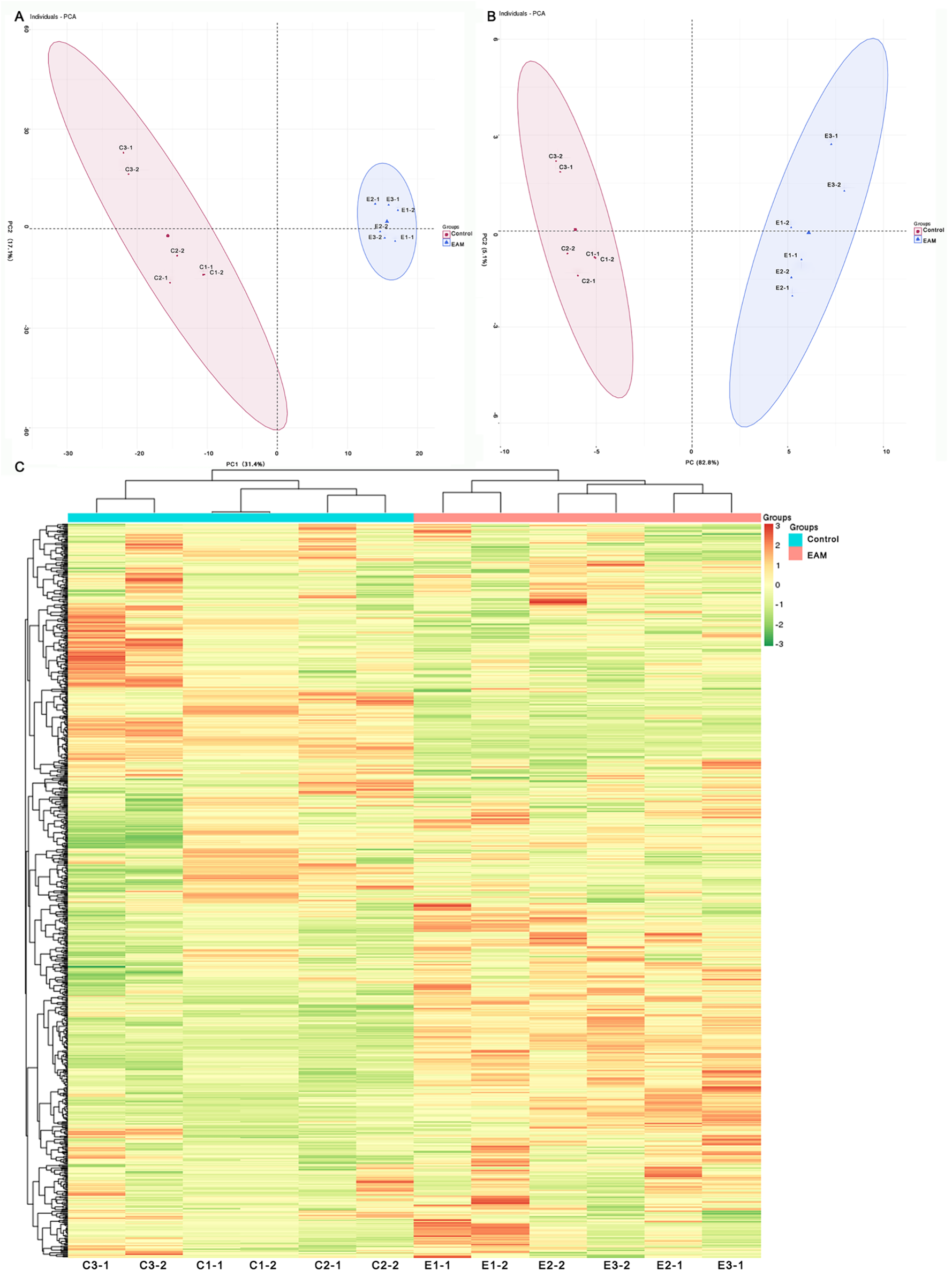
Statistical analysis of the urine proteome of EAM rats. A) PCA of the 816 proteins from the EAM and control urine proteomes on day 21. Each axis is labeled with the variation in the data. B) PCA of the 46 changed proteins between the EAM and control urine proteomes. C) Hierarchical clustering of the 816 proteins from the 12 samples (six subjects in the EAM and control groups) at day 21. Lines represent proteins, and the colors correlate with their abundance (red indicates more abundant; green indicates less abundant).

In addition to determining individual differences, differences between technical replicates and the effects of myocarditis on the complete proteome, all quantitated proteins were also investigated by principal component analysis (PCA). Twelve quantitative datasets were exported from the Scaffold software (including six day-21 samples from the EAM and control groups and the replicate of each MS analysis) and were included in the PCA. The biplot shows the clustering and trends present in the protein profiles. Samples that have similar scores tended to form clusters, whereas samples with dissimilar scores were at greater distances apart (Figure 3A). The quantitative urine proteome representations for day 21 in the EAM group are different from those in the control group since proteome representations do not overlap.

As the urine proteome of the EAM and control groups are quite different, the changes in proteins were then calculated. The changed urinary proteins were defined as those with a p value < 0.05 by two-sided, unpaired t-test and a fold change > 1.5 compared with controls. Among all of the identified proteins, 5.6% met the strict criteria, and among them, 32 proteins were significantly increased 21 days after myosin immunization, while 14 proteins were significantly decreased. The molecular weight of these changed proteins ranged from 10 kDa to 190 kDa. Detailed information on these changed proteins in EAM rat urine is provided in Table 1. PCA was also conducted for the 46 changed proteins, and the changed proteins between the EAM and control groups also formed two clusters (Figure 3B).

### Changed proteins previously reported in serum or cardiac tissue from patients with myocarditis

Changed proteins that are homologous to humans are potentially useful in clinical practice. By importing the changed proteins into InParanoid [22], the protein orthologs can be viewed. Among them, 40 out of 46 proteins had human orthologs, and they may likely be investigated in human myocarditis in future studies.

Among the 40 proteins that have human orthologs, ten were previously reported in serum or cardiac tissue from patients with myocarditis or cardiac injury (Table 1), which may indicate the accuracy of the MS analysis and show that urine is a good source of myocarditis biomarkers. Several examples to illustrate the performance of these proteins in urine and serum/cardiac tissue are as follows:

Regenerating islet-derived protein 3 (REG3) is involved in acute phase signaling. Reg3beta was considered to be an important regulator of macrophage trafficking to the damaged heart, and the macrophages are essential for cardiac healing after myocardial ischemia[23];

Cystathionine gamma-lyase (CSE) is an enzyme that markedly reduces H_2_S levels in the serum, heart, aorta and other tissues. Several studies have found that serum CSE mRNA and CSE protein levels are increased in coxsackievirus B3-induced myocarditis[24];

Myoglobin is an intracellular oxygen-transport protein. Myoglobin protein expression in heart tissue may reflect the intensity of myocarditis and serve as one of the pathogenetic factors of cardiac dysfunction in myocarditis[25];

Matrix metalloproteinases (MMPs) are regulators of the extracellular matrix. After coxsackievirus B3 infection, extracellular matrix remodeling is triggered, and this response may involve the activation of MMPs[26];

After the onset of acute myocardial infarction, heart fatty acid-binding protein (H-FABP) within cells is released into plasma. As a result, cytoplasmic H-FABP is suitable as an early indicator of acute myocardial infarction[27];

ATP5b was significantly increased mouse heart tissues during the acute phase of coxsackievirus B-induced myocarditis and was significantly decreased during the chronic phase[28];

Beta 2-microglobulin is considered to be a useful index for monitoring the development of myocardial inflammation[29];

E-selection plays an important role in inflammation, and increased cardiac expression of E-selectin has been demonstrated in mice with myocarditis [30]; and Changes in plasma lactate dehydrogenase activity have also been reported in myocarditis patients[31].

Interestingly, the ten proteins that were previously reported to be related to myocarditis were not the most significantly changed proteins in this study, which may indicate that additional proteins may be candidate biomarkers of cardiac injury in the future.

### Western blot analysis

Four changed proteins (myoglobin, cystathionine gamma-lyase, l-lactate dehydrogenase and heart fatty acid-binding protein) that have human orthologs, have commercially available antibodies and were not previously shown to be increased or decreased in urine from myocarditis patients were verified by western blot at three-time points (baseline, day 14 and day 21) in twelve additional EAM rats (Figure 4).

**Figure 4.**
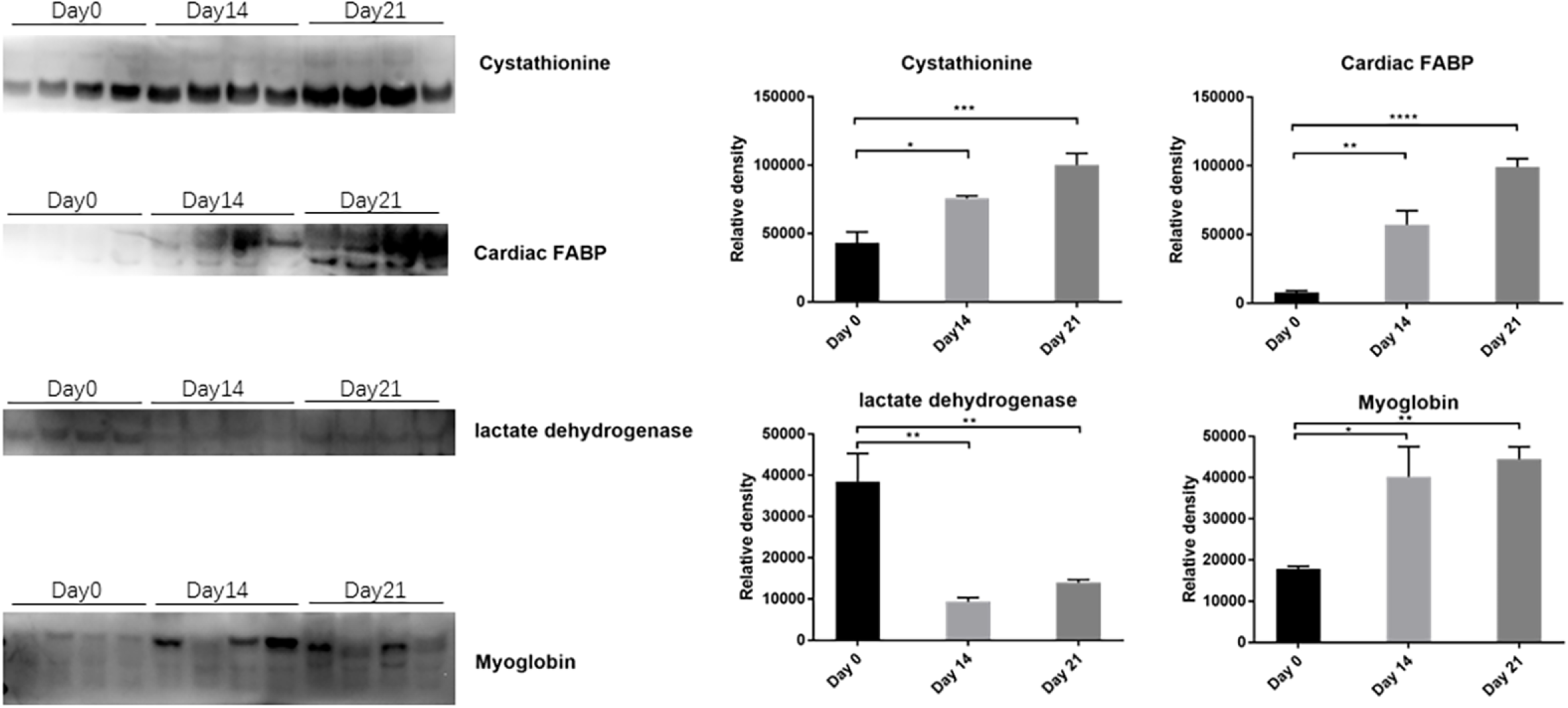
Western blot was used to validate four changed proteins in twelve additional rats. The proteins were regenerating islet-derived protein 3, myoglobin, cystathionine gamma-lyase and beta 2-microglobulin. * indicated a p value <0.05, ** indicated a p value <0.01, and *** indicated a p value <0.001.

Urine cystathionine gamma-lyase (CSE) levels in the EAM group on day 14 were over 1.5-fold greater than those in the control group and continued to increase thereafter. On day 21, the urinary CSE levels were 2.3-fold higher in EAM rats than in controls. The l-lactate dehydrogenase protein was down-regulated in the urine of EAM rats on day 14 and day 21. On day 14, the protein level of l-lactate dehydrogenase was markedly decreased (p value < 0.01) in EAM rats compared to control rats and remained low on day 21. The urine of EAM rats exhibited a higher level of myoglobin compared with that of control rats (p<0.05), but the changes were not statistically significant on days 14 and 21. Protein levels of l-lactate dehydrogenase and myosin were significantly different between EAM and control rats from baseline to day 14 and remained at a similar level on day 21. The western blot results of the three above mentioned proteins were consistent with the shotgun proteomics data.

However, in this study, urinary heart fatty acid-binding protein (H-FASP) was down-regulated in EAM rats, as identified by MS. In this case, the western blot results were not consistent with the MS results. Urinary H-FASP levels were significantly increased (p value <0.01) during the occurrence and progression of myocarditis. This may partially be because the proteins identified by MS were quantitated by summing all fragments, whereas the western blot experiment focused only on the protein bands with a molecular weight consistent with the candidate protein. When the trends of protein expression during myocarditis are analyzed in the future, the methods of diagnosis should be seriously considered.

### Twelve genes were annotated in cardiovascular functions by IPA

To assess the involvement of dysregulated genes in genetic networks, IPA software was used to reveal their association with important biological functions. One of the most significant gene networks that was associated with dysregulated genes in the EAM samples was cardiovascular function. A total of 12 changed proteins identified in the present work were involved in cardiovascular function (Figure 5), which may indicate the high accuracy and reliability of this study. The molecules involved in the cardiovascular function network are commonly associated with cardiac dysfunction, such as the expression of collagen. Additionally, the IPA highlighted MMP2 and COL1A1, which were both determined to be core components related to other proteins in the connectivity map generated in the current study. MMP2 mRNA expression in the human heart is associated with deteriorating dilated cardiomyopathy[32], and myocarditis may lead to dilated cardiomyopathy if left untreated. In addition, COL1A1 mRNA is expressed in primary microvascular endothelial cells from the human heart[33]. Both MMP2 and COL1A1, which are closely related to heart function, were changed in EAM rat urine. Nevertheless, the gene products identified in the EAM study that are not involved in the cardiovascular function network may also play important roles in myocarditis disease progression and contribute to elucidating the mechanism of myocarditis.

**Figure 5.**
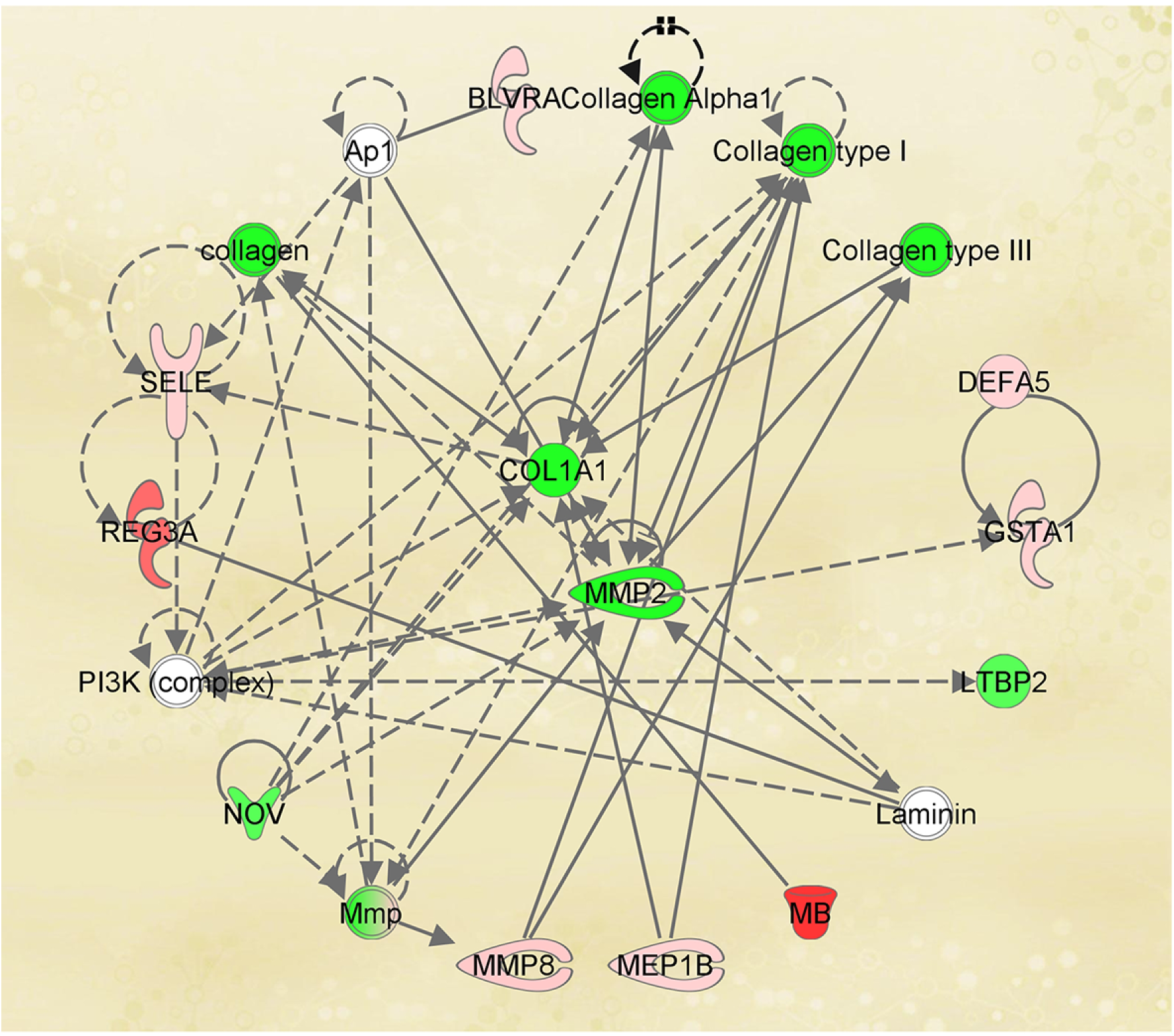
IPA determined that the changed proteins are involved in the cardiovascular network. Red indicated up-regulated genes, and green indicated down-regulated genes in the present study.

### Functional analysis of all changed proteins

To test the specificity of these selected urinary biomarkers for the detection of myocarditis, we compared our results with a previous manually curated urinary biomarker database[34]. Thirteen proteins have been previously identified as urinary candidate protein biomarkers for different diseases. Among those, eight proteins exhibited the same trend in other diseases and as in the EAM rat model; for example, the Ig gamma-2A chain was up-regulated in the urine of renal calculi patients and recurrent renal calculi patients[35], and latent-transforming growth factor beta-binding protein 2 was significantly elevated in both the serum and urine of patients with Kawasaki disease[36]. Five proteins that were decreased in the urine of EAM rats have been reported to be increased in the urine of other diseases in previous studies. Fatty acid-binding protein (heart) was significantly elevated in normoalbuminuric diabetic patients[37], whereas an opposite trend was observed in EAM urine.

To further explore the characteristics of the changed urinary proteins during EAM development, Gene Ontology (GO) analysis was performed to provide insight into the most common cellular components and molecular functions. Approximately 40 out of 46 changed proteins were annotated in the GO molecular function analysis, and ten proteins, such as regenerating islet-derived protein 3, E-selectin and beta-2-microglobulin, were enriched in immune system processes that play important roles in immune myocarditis development. In addition, the changed proteins were involved in metabolic processes (37%), localization (25%) and cellular processes (23%). Although proteins from all parts of the cell could be detected, extracellular matrix proteins were significantly over-represented in the main cellular component.

## Conclusion

In this study, candidate urinary biomarkers of myocarditis were uncovered in EAM-induced rat models. Many of them had previously been reported to be related to myocarditis. Additionally, about one-third of the corresponding differential genes were annotated to participate in the cardiovascular function network. In conclusion, the study showed that urine can be a good source of myocarditis biomarkers.

## Author Contributions

M.Z. and Y.G. prepared the first draft. M.Z., X.L. and Y.G. conceived and designed the experiments. M.Z., X.L. and J.W. performed the experiments. M.Z. and J.W. analyzed the data. All authors approved the final manuscript.

## Acknowledgements

This work was supported by National Key Research and Development Program of China (2016YFC1306300), National Basic Research Program of China (2013CB530805), Beijing Natural Science Foundation (7173264, 7172076), the Fundamental Research Funds for the Central Universities (2015KJJCB21), Beijing cooperative construction project (110651103) Beijing Normal University (11100704).

---

**Supplemental Table 1. All identified urinary proteins in this study.**

